# Drug Response Modeling across Cancers: Proteomics vs. Transcriptomics

**DOI:** 10.1101/2024.12.04.626700

**Authors:** Zetian Zheng, Lei Huang, Fuzhou Wang, Linjing Liu, Jixiang Yu, Weidun Xie, Xingjian Chen, Xiangtao Li, Ka-Chun Wong

## Abstract

Cancer cell lines are the most common *in-vitro* models for the evaluation of anti-cancer drug sensitivities. Past studies have been conducted to decipher and characterize the pharmacogenomic feature of cell lines based on other omics data, such as genomic mutation data and whole-genome RNA sequencing (RNA-seq) profiles. In particular, proteomic data is also an essential component for the characterization of tumours. However, different from RNA-seq datasets rich in numerous transcriptome profiles of cancer cell lines and cell viability assay of drug responses, the pharmacogenomic protein quantifications are relatively scarce. With the availability of the recently enriched proteomic dataset ProCan-DepMapSanger, we systematically evaluated the interplays among genomic mutations, transcription, and protein expressions across cancer cell lines. In general, blood cancers have higher RNA-protein correlations than those in solid cancers. The differential expression analysis on protein data helped identify more expressional and functional impact of genomic mutations of cancer genes. We also integrated the proteomic map with drug molecular chemical features to construct a bi-modal machine learning model to infer the drug sensitivities of cancer cell lines. Our results demonstrated that protein quantifications can lead to better drug response prediction performance than the model trained on transcriptome profiles. In addition, integrating protein data with drug chemical features, represented as molecular graphs and learned by Graph Neural Network, outperformed the state-of-the-art model DeepOmicNet for drug response prediction in proteomics.

## Introduction

The heterogeneity of drug response phenotypes challenges the clinical implementation of anti-cancer drugs^1^. Many researches have been conducted based on the pharmacogenomic information of cancer cell lines from public databases (e.g., GDSC, CCLE)^2,3^. Those studies aim to discover the putative therapeutic biomarkers to understand the perturbational consequences as well as refine the personalized treatment recommendations^1^. Pertaining the same object, many efforts have also been made by the computational community to construct machine learning (ML) models to assist drug sensitivity predictions, most of which were built based on gene expression profiles^4^ or genomic signatures^5^. Previous studies reported that the gene-level RNA-protein Pearson correlations among tumors are approximately 0.5 in many cancer types, such as HPV-negative head and neck squamous cell carcinoma (HNSCC)^6^ and lung adenocarcinoma (LUAD)^7^. The correlation coefficients are even lower than 0.5 in other cancer types, such as pancreatic ductal adenocarcinoma (PDAC)^8^ and clear cell renal cell carcinoma (ccRCC)^9^. This discrepancy suggests that RNA expression and protein quantification are not strongly correlated at the genome-wide level. An emerging consensus has indicated that this phenomenon is influenced by the combination of multiple mechanisms, such as signaling pathways, genomic alterations, and post-translational modifications of proteins^8^. Therefore, it is suggested that the large-scale proteomic data may differ from transcriptomic data and should be evaluated separately.

The incompatibility between RNA and protein expression levels subtly affects the effectiveness of anti-cancer treatments. It is widely believed that gene expression profiles and the gene alterations of tumours alone are insufficient to explicate the drug response heterogeneity and provide progress for precision medicine^8^. Albeit the genomic mutational analysis and transcriptomic sequencing profilings could improve the characterization of tumours for the subclass stratification of patients, prognosis prediction, and clinical treatment recommendations^10^, neither of them can accurately predict the changes in the corresponding protein expression levels and functional status^11^. In contrast, researchers revealed that proteomics could better unveil the downstream influence of copy number alterations and somatic mutations that are not evident in transcriptomic profiles^12^. Moreover, although RNA profiles could be applied to identify specific pathways that are considered to be active in patient subgroups, the anti-cancer treatments that target these pathways usually fail^12^. One possible explanation for the clinical translation failure in targeting these pathways is that the molecular mechanisms identified by RNA transcriptomes exit in many regulatory layers beyond the proteins^12,13^. Indeed, the basic functional units of biological processes of cells are proteins, which are the primary targets of anti-cancer treatments^14^. More specifically, the anti-cancer compounds help control the circuitry of complex cellular networks and influence the activities of protein workforce^15^. The limitations described above hinder the clinical translation of findings based on genomic sequencing techniques in the absence of proteomic profiles^8^. Overall, proteomics could provide us with new insights into perturbational biology, which differs from transcriptomics and genomics data types^16^, enabling us to identify the putative therapeutic targets of anti-cancer drugs^8^.

There is a pressing need to evaluate the inference capability of drug response and cancer vulnerability based on proteomics on a larger scale, both at a genome-wide level and across various types of cancer. In light of this, an open-source pan-cancer cell line proteomic map was recently established in ProCan-DepMapSanger database, quantifying more than 8,000 proteins across 949 cancer cell lines^16^. This database provides a valuable resource of proteomic data and a constructive reference for drug-proteomic analysis. Existing pharmacogenomic analysis mainly relies on genomics and transcriptomics. It is now possible to evaluate drug sensitivity and refine the pharmacogenomic interactions in such models by introducing protein data^16,17^. The ProCan-DepMapSanger dataset contributors also built a deep learning model (DeepOmicNet) to evaluate the predictive power for drug sensitivity inference on cancer cell lines based on proteomic and other omics datasets^16^. Unfortunately, we noticed two limitations of DeepOmitNet, as described below.

Firstly, the number of output layer neurons is the number of evaluated drugs (n = 625)^16^. This kind of dataset-specific neural network architecture makes the model less generalizable. For instance, to accommodate user requests for customized analyzed drugs, it is necessary to rewrite and retrain the entire model framework accordingly, especially when additional drugs are not included in the previously evaluated drug list. This is because DeepOmicNet only takes a single type of omics data as input during the training process. This drawback makes DeepOmicNet vulnerable to predicting the drug response efficacy of novel drugs^16^. There are also many multi-modal machine learning models that use both gene expression (transcriptome) and drug molecular chemical features (represented as a graph) as inputs^18^. However, due to the lack of a protein quantification dataset, this kind of bi-modal ML framework has not been widely used in large-scale proteomic data. In addition, due to the evident heterogeneity between RNA transcript expression and protein quantification, the drug response prediction ability of bi-modal ML model that considers both large-scale proteomics and drug molecular chemical features has not been widely evaluated. Therefore, we utilized a Bi-modal Drug Response Network (BDRN) that integrates proteomics with drug chemical features (using graph neural network) to evaluate the drug response prediction capability of proteins and compared it with that of RNAs.

Secondly, during prioritizing drug responses for the promising candidate drug selections, the authors did not consider the differences between hematological malignancies and solid cancers^16^. However, it was believed that hematological malignancies result from abnormal hematopoietic stem cell differentiation and have a distinct cancerogenesis origin from solid cancers^19^. Moreover, previous studies demonstrate that hematological malignancies and solid cancers show different disease trajectories, side effects of anti-cancer therapy, and heterogeneous treatment outcomes. Therefore, they need specific treatment paradigms accordingly^20^. For instance, both Chimeric antigen receptor (CAR)-modified T (CAR-T) cells and Bispecific T cell engager (BiTE) immunotherapies demonstrated promising anti-cancer efficacy in B cell hematological malignancies^21^. However, the efficacy of treatments for solid cancers fell short of expectations in the early phase clinical trials, indicating a need to conduct further research and treatment development for the above immunotherapies on solid cancers^21^. Therefore, we comprehensively characterize cancer cell line transcriptomics and proteomic data. Our results demonstrated that, in general, the RNA-protein correlation coefficients in hematological malignancies are significantly higher than those observed in solid cancers (Fig. 2b, Fig. S4). Moreover, the anti-cancer drugs that responded differently between hematological malignancies and solid cancers were also identified. For instance, we found that the overall predictive power (ground-true vs. predicted ln(IC50) Pearson coefficient) for drug Volasertib in hematologic maligancies was only around 0.34, while it was around 0.66 in solid cancer cell lines (Fig. 5).

**Figure 1.**
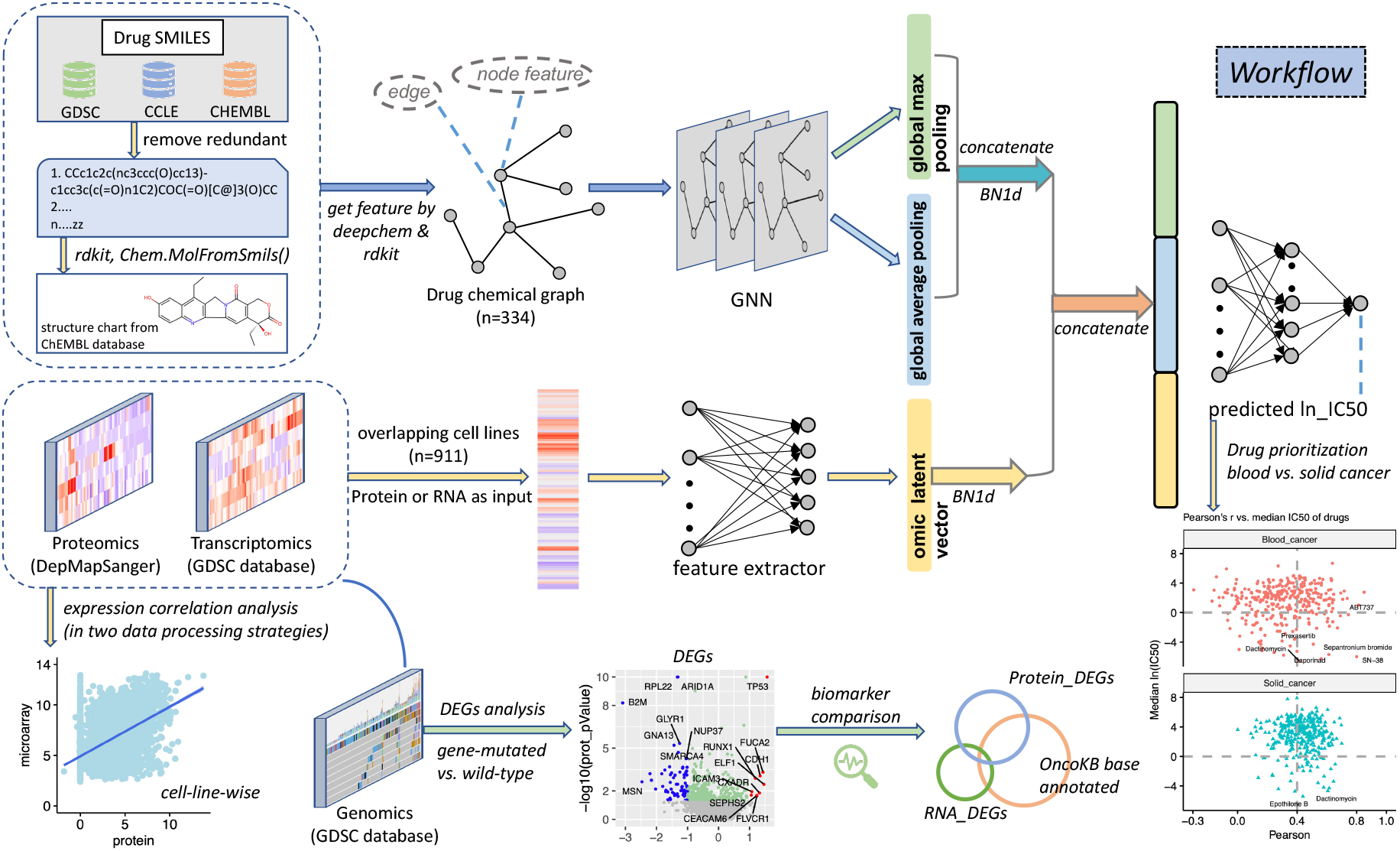
Workflow of bi-modal Graph Neural Network for drug response prediction based on proteomics or transcriptomics. The workflow of BDRN for drug response prediction is on drug chemical features (represented as a graph) and the expression levels of protein (or RNA). Drug chemical features were processed and obtained through deepchem^25^ and rdkit packages^26^. The drug chemical feature was learned via graph neural network^27^, and a global max pooling and a global mean pooling were applied to learn the feature map of each graph. Meanwhile, MLP layers were used as feature extractors to obtain the latent vector of RNA and protein data. The normalized latent vectors of graph and protein (or RNA) were concatenated, and the concatenated feature map for drug and omic features was fed into the predictor for the inference of drug responses. The detail for data collection and data preprocessing was described in the Method section. Outputs of BDRN (predicted ln(IC50)s) were applied for drug response sensitivity prioritization. We analyzed hematological cancers and solid cancers separately due to an evident discrepancy between them. Moreover, we conducted cell-line-wise RNA-protein expression correlation analysis. Next, the interplays among genomic mutations, transcription, and protein expressions across cancer cell lines were systematically evaluated. The candidate biomarkers (either reflected in RNA and protein level) were compared with oncogenesis-related cancer genes and druggable genes that were annotated in the precision oncology base OncoKB to evaluate their functional impacts.

**Figure 2.**
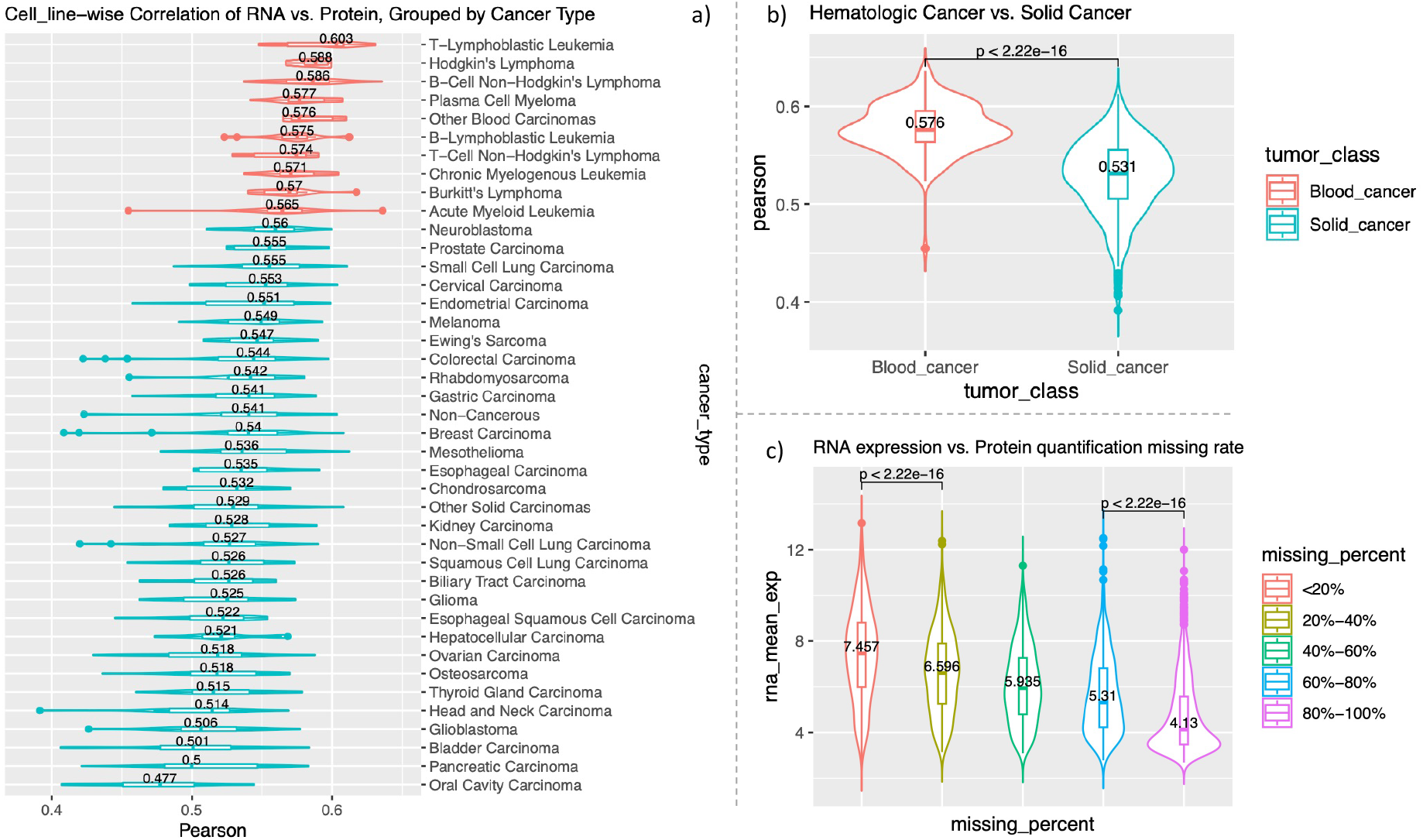
Pan-cancer cell line RNA-protein correlation landscape. a) Boxplot with violin plot for the cell line-wise correlation between RNA and protein expression level; missing values of proteins were replaced as zeros. Cell lines were grouped by cancer types and sorted by the values of Pearson’s r. b) Boxplot for comparing RNA-protein cell line wise correlation coefficients between hematological cancers and solid cancers. c) Boxplot for the RNA expression level regarding the genes with different protein quantification missing rates; genes were assigned into five groups based on the missing rate of protein quantification data in all cancer cell line populations.

In addition, we have conducted a detailed comparison between RNA sequencing data and protein sequencing data. The current data-independent acquisition mass spectrometry (DIA-MS) based proteomic data contain a large proportion of missing values^16^ due to both technical and biological aspects. The data processing method for missing values will greatly impact the analytical and statistical results of downstream analysis^22^. We believe that this drawback to some extent can be mitigated by referring to the expression profile of the matched RNA expression levels in the context of the central dogma^23^. To address this, we systematically compared the difference of results for the RNA-protein Pearson correlation and differential expression analysis (both at RNA and protein quantification levels) when missing values were directly removed during the correlation analysis and that when missing values were treated as zeros. We found that removing all missing values will lose plenty of expression information at both RNA and protein levels during the correlation analysis (Fig. S5). In addition, we observed that genes with higher missing rates of protein quantification data also exhibited significantly lower RNA expression levels.

In contrast, replacing missing values as zeros not only obtained higher RNA-protein expressional coefficients than removing all missing values; it also enables the identification of additional differentially expressed biomarkers (either at RNA or protein level) and carcinogenesis-related genes that overlapped with cancer genes that annotated in OncoKB database (https://www.oncokb.org/)^24^. More specifically, two druggable genes ARID1A and TP53 were also found to be up-regulated at protein levels among gene-mutated cell lines, compared to wild-type ones. In contrast, no apparent expression fluctuations were observed in RNA expression levels for the above two druggable genes. Our results suggested that the protein map is a fundamental component for the evaluation of the expressional and functional impact of genomic mutations. It is also necessary to integrate a proteomic map to explore potential druggable targets.

## Results

### Prediction performance of our proposed GNN models and baseline model DeepOmitNet

Prior research has indicated that the RNA-protein Pearson correlation in tumors is approximately 0.5^7^ or lower^9^. Moreover, there remains a gap in our understanding on the correlation between RNA and protein expression at pan-cancer scale. To address this problem, we systematically conducted correlation analyses at both gene-wise level and cell-line-wise level. We observed that the median gene-wise Pearson correlation between RNA expression and protein quantification is around 0.436, while the median cell-line-wise RNA-protein Pearson correlation is 0.539 (Fig. S2). This result suggested that genes may have varying impacts on how a cancer cell line responds to drug perturbations at the levels of RNA transcripts and proteins. Therefore, we compared the machine learning performance between RNA and protein data at the level of cell line.

Furthermore, to address the limited scalability of the previous DeepOmitNet model^16^, we have also incorporated the compound chemical feature (represented as a graph) of drugs. Although integrating drug graph features with RNA expression data of cancer cell lines is a commonly used approach for predicting drug response, the potential benefits of incorporating drug graph features in large-scale proteomic analysis of cancer cell lines at the pan-cancer level have not been widely explored in terms of the drug response prediction capability. Therefore, we utilized a framework to extract molecular graph features of drugs to predict the corresponding responses. More specifically, we applied separate feature extractors for transcriptomic and proteomic data. The number of nodes in the first layer of the feature extractor matches with the number of measured genes in the RNA and protein data, respectively (see Method). Next, two different graph convolutional layers (GCNConv and SAGEConv from PyTorch Geometric^27^) were adopted to learn and extract the compound chemical features of drugs. We conducted a 5-fold cross-validation (CV) strategy for hyper-parameter selection based MSE loss metric on each fold’s validation set. To ensure a fair comparison, we utilized the same metric in previous study^16^, which calculates the cell line-wise Pearson’s r for each drug individually. We averaged the predicted ln(IC50)s from the five models with minimal validation loss at each fold. The drug response prediction performance on the cell lines of the held-out test set for different omics data types (RNA and protein) and different convolutional layers were summarized in Table 1 and Fig. S1. Compared to RNA data in the test set, the BDRN models demonstrate slightly better performance (cell line-wise person’s r of ground-true ln(IC50) values vs. predicted ln(IC50) values for each drug) on the protein data in the test set with both GCNConv (protein: 0.5467 vs. RNA: 0.5311) and SAGEConv (protein: 0.5463 vs. 0.5272) layers (Fig. 1, Table 1, Fig. S1). Moreover, all four bi-modal models obtained higher Pearson’s r than the baseline model (DeepOmitNet), whether comparing the median (0.5261) or the mean Pearson’s r (0.5058, Table 1, Fig. S1). Our bi-modal model results indicate that the proteomic data can obtain comparable (and slightly better) drug response prediction performance than transcriptomic data. Furthermore, this model learns the representation of chemical features of drugs (represented as graphs), which is suitable for predicting the drug response efficacy of novel drugs (unseen in the training process) containing learned drug chemical features^16^, therefore has better scalability than DeepOmicNet. Our result suggested that proteomic data may offer us new insights for analyzing anti-cancer drug treatments. In addition, based on the previous reports that RNA-protein expressions are not strongly correlated^6–9^, the biomarkers that associate with drug responses may differ between RNA and protein levels. Therefore, it is necessary to systematically compare the similarities and differences in RNA and protein expression levels.

**Table 1.**
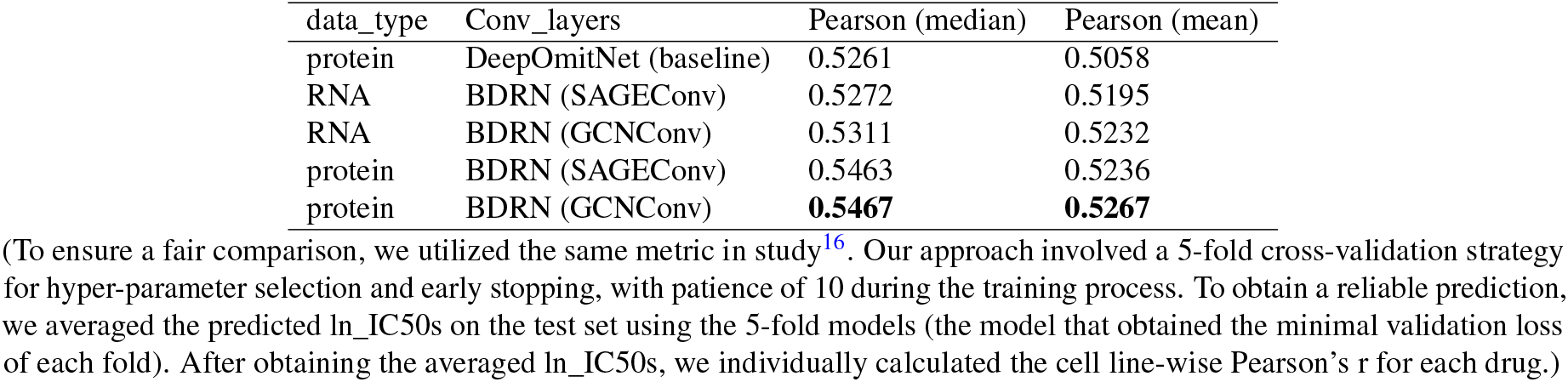
Prediction performance of our proposed GNN models and baseline model DeepOmitNet

### Pan-cancer cell line RNA-protein correlation landscape

The currently released ProCan-DepMapSanger database quantifies over 8,000 proteins across 949 cancer cell lines^16^, enabling us to re-analyze the association among genomic mutations, transcriptomic profiling, and proteomic quantifications of tumours in more details. This enriched pharmacogenomic resource allows us to refine the characterization of cancer heterogeneity and anti-cancer biomarker discovery. We systematically evaluated the difference in RNA-protein expression coefficients across different cancer types of cell lines. The association of the expression levels of RNAs and the missing rate of corresponding protein levels was also explored by grouping genes into five groups based on the missing rate of protein data. Next, we conducted the differential expression analysis at both RNA and protein expression level based on the mutation status of genes across the cell line population. To find evidence-based promising biomarker genes, we integrated differential expression genes (DEGs) with the cancer-gene list and druggable-gene list annotated in OncoKB database.

#### In general, hematological malignancies possess higher RNA-protein Pearson correlation than solid cancers

As described in the introduction section, there is an evident discrepancy between RNA transcript and protein quantification levels. In order to capture a more refined pan-cancer RNA-protein correlation landscape, we grouped cancer cell lines by annotated cancer type summarized in previous study^16^ and calculated the RNA-protein correlation for each cancer type, respectively. One challenge for DIA-MS-based proteomic data is the missing values^16^. The existence of missing values may be due to numerous biochemical mechanisms, such as miscleavage, ionization competition^28^. It may also originate from bioinformatic disadvantages, such as peptide misidentification, and precursors ambiguous matching^28^. To investigate what extent the missing values will impact the RNA-protein expression correlation analysis results, we analyzed two different scenarios: one involved deleting all missing values (and the corresponding RNA expression data of genes with missing protein quantification values were also ignored during analysis), while the other replacing all missing values as zeros.

For the first strategy, all missing values in the proteomic data were excluded when calculating the gene-wise and cell-line-wise correlation coefficients. Under this scenario, the Pearson correlation between RNA and protein expression is strongest in T-Lymphoblastic Leukemia (Median Pearson’s r = 0.401) and weakest in Oral Cavity Carcinoma (Median Pearson’s r = 0.316, Fig. S3). In general, hematological malignancy cell lines exhibited higher rankings than solid cancer ones when sorting cell lines by Pearson’s r of RNA-protein correlation coefficients. Specifically, the top-four cancer types exhibiting the highest correlation coefficients all fall within the category of hematological malignancies, whereas the bottom-four cancer types with the smallest correlation coefficients all belong to solid cancers (Fig. S3). Upon classifying the cell lines into blood cancers and solid tumors, we observed a significant difference in RNA-protein Pearson correlation coefficients between these two cancer groups, with blood cancers showing significantly higher correlation coefficients (Median Pearson’s r = 0.389 vs. Median Pearson’s r = 0.371, p<0.001, Fig. S4).

Next, we followed the processing strategy used in DeepOmicNet^16^, which involved replacing all missing values with zeros. Interestingly, the RNA-protein expression’s Pearson correlation coefficients for all cancer types were increased. In addition, all hematological malignancy cell lines obtained higher RNA-protein correlations than solid cancers after ranking by Pearson coefficients (Fig. 2a). Similar to above result that excluding missing values, the correlation of RNA-protein is strongest in T-Lymphoblastic Leukemia (Median Pearson’s r = 0.603) and weakest in Oral Cavity Carcinoma (Median Pearson’s r = 0.477, Fig. 2) when missing values were replaced with zeros. Likewise, the hematological malignancies exhibited significantly higher RNA-protein Pearson correlations than that in solid tumors (Fig. 2b, Median Pearson’s r = 0.576 vs. Median Pearson’s r = 0.531, p<0.001). Our results suggested that the Pearson correlation coefficients between RNA and protein transcription levels increased when missing values were treated as zeros. In general, hematological malignancies exhibited higher RNA-protein correlation than solid tumors, which was consistently observed across both data processing methods.

#### RNA expression levels decrease when protein quantification missing percentage increase

To find out the possible biological mechanisms that could explain why replacing missing values with zeros led to increased absolute values of RNA-protein Pearson correlation coefficients compared to excluding all missing values, we conducted gene-wise RNA expression comparisons based on the missing rate of the corresponding proteins. Firstly, we calculated the missing rate of protein values across all cancer cell lines for each protein-coding gene. We then assigned each gene into five groups based on the missing rate of the corresponding protein (Fig. 2c). We found that the gene group with the smallest missing rate (<20%) of protein levels obtained the highest RNA expression level and the largest RNA expression variance (variance = 4.13) among all five groups, while the gene group with the highest missing rate (80%-100%) of protein levels obtained the lowest RNA expression level, as well as the lowest RNA expression variance (variance = 2.62, Fig. 2c).

To better visualize the results, we have also carefully compared the difference between these two missing data processing strategies by a scatter plot, taking glioblastoma cell line GAMG as an example (Fig. S5). Our analysis revealed a peak in the density plot within the range of 2 to 3 (RMA-normalized) low RNA expression levels when considering the entire RNA profile (only genes included in both transcriptomic data and proteomic dataset were included). In addition, a similar peak in the density plot within the range of 0 to 1 (DIA-NN processed^29^) low protein expression levels was found when protein missing values were treated as zeros; the RNA-protein expression correlation Pearson’r is 0.52 in the scenario (Fig. S5a). On the contrary, the RNA expression data within the peak of 2 to 3 (RMA-normalized) were not included for the calculation of RNA-protein correlation when protein missing values were excluded; and Pearson’s r dropped to 0.37 in this processing strategy (Fig. S5b). This may be attributed to the exclusion of genes with missing proteins, which results in the loss of information for a large proportion of low-expression RNAs and low-expression proteins. As a consequence, the slope of the RNA-protein Pearson correlation coefficient regression line decreased (Fig. S5a, Fig. S5b).

Our results indicated that diverse biological mechanisms for cancer carcinogenesis and tumor progression may exist between hematological malignancies and solid cancers. In fact, the heterogeneity between these two types of cancer is not only limited to differences in the correlation of RNA expression and protein quantification levels but also manifested in differences in the treatment and prognosis of cancer patients in the real world. It was suggested that the tumor heterogeneity between hematological malignancies and solid cancers leads to diverse disease trajectories and requires specific treatment paradigms^20^; patients with hematological malignancies often suffer significant toxicities and side effects on clinical anti-cancer medications^20^. Therefore, based on the discrepancy between these two types of cancer, it is necessary to treat them separately when conducting candidate drug screening and downstream promising drug selection analysis.

#### Genetic mutations affect the expression changes of RNA and proteins to varying degrees

It was suggested that the genomic mutation map together with transcriptomic profiling could improve the stratification of cancer patients, and refine the clinical treatment recommendation^10^. However, neither of them could reflect the fluctuations in protein expressions^11^. Thanks to the recently released Pan-cancer cell line ProCan-DepMapSanger proteomics dataset^16^, the transcriptomic dataset processed by authors of previous research^4^, and the genomic mutations of GDSC cancer cell line database that summarized in previous study^17^, we were able to evaluate the effect of genomic mutations on the fluctuations in both RNA expression and protein expression levels at Pan-cancer scale. For each gene, the two-sided Wilcoxon-test was conducted to compare RNA and protein expression levels between gene-mutated cell lines and the gene wild-type cell lines. In addition, to ensure more reliable statistical comparison results, we required that each gene being compared must simultaneously have at least 10 mutated cell lines and 10 non-mutated cell lines. To further compare the differences between the data processing strategies mentioned above, we continue our discussion of both approaches in this section.

Firstly, all missing values in the ProCan-DepMapSanger proteomics dataset were excluded in the Wilcoxon rank sum test calculating step. Under this scenario, we observed that only a small number of genes exhibited differential expression between the mutated and non-mutated groups, regardless of whether the tests were conducted at the RNA (Fig. S6a) or protein level (Fig. S6b). We believe that this may be due to a significant loss of quantification data resulting from the sample size requirement for each group to have at least 10 cell lines and the removal of genes with missing values, which leads to a notable reduction in the number of genes used for comparison (Fig. S5b).

Secondly, we replaced all missing values with zeros. Adopting this processing strategy can preserve all measured values of RNA expression data for all cancer cell lines. Compared to the previous processing method, we observed a significantly higher number of RNA differentially expressed genes (Fig. 3b-c, Fig. S6a-b). Moreover, we observed a comparable trend in protein-level differential expression to that seen in RNA level, where most genetic mutations resulted in decreased protein expression levels, with only a few genes exhibiting increased protein expression levels after the mutation (Fig. 3c). The convergence towards a similar trend suggests that this data processing approach is valid. In addition, compared to the wild-type cell lines, both RNA and protein expression levels of three genes (CDH1, CXADR, FLVCR1) showed significant increment in the mutated cell lines. Compared to the wild-type cell lines, nine genes (B2M, DPYSL3, GSN, MAP1B, MSN, NNMT, RAB34, SPARC, TPBG) presented a significant decrease in the mutated cell lines at both RNA and protein expression levels (Fig. 3b,c). The mutation landscape of genes that exhibited either up-regulated or down-regulated expression in either RNA or protein levels were presented in Fig. 3a (this figure includes only those genes with a mutation frequency of at least 3% of whole cell line population). No matter in hematological malignancies or solid tumor cell lines, the most frequently mutated gene is TP53 (68% in total, Fig. 3a), and the mutated cell line group exhibits significantly higher protein expression levels compared to the TP53 wild-type group (Fig. 3c). Two genes (ARID1A, SMARCA4), with mutation frequencies over 10% of the whole cell line population, showed a significant decrease in protein expression levels in their corresponding mutated cell lines compared to their wild-type cell lines. (Fig. 3a, Fig. 3c). Compared to wild-type cell lines, we further observed several genes that were consistently up-or down-regulated in both RNA and protein expression levels in the mutated cell lines, such as gene FLVCR1 and MSN (Fig. 3b-c). Additionally, we found that some genes exhibited significant fluctuations at the RNA level but non-obvious changes at the protein level due to gene mutations; while others showed evident changes at the protein level but are not reflected in RNA abundance (Fig. 3b-c). Our results revealed that there may be significant differences in the impact of genetic mutations on RNA and protein expression levels among different genes, and the changes in RNA levels due to genetic mutations may not be able to reflect the expression fluctuations of proteins, which highlights the essential complementary role of proteomic data.

**Figure 3.**
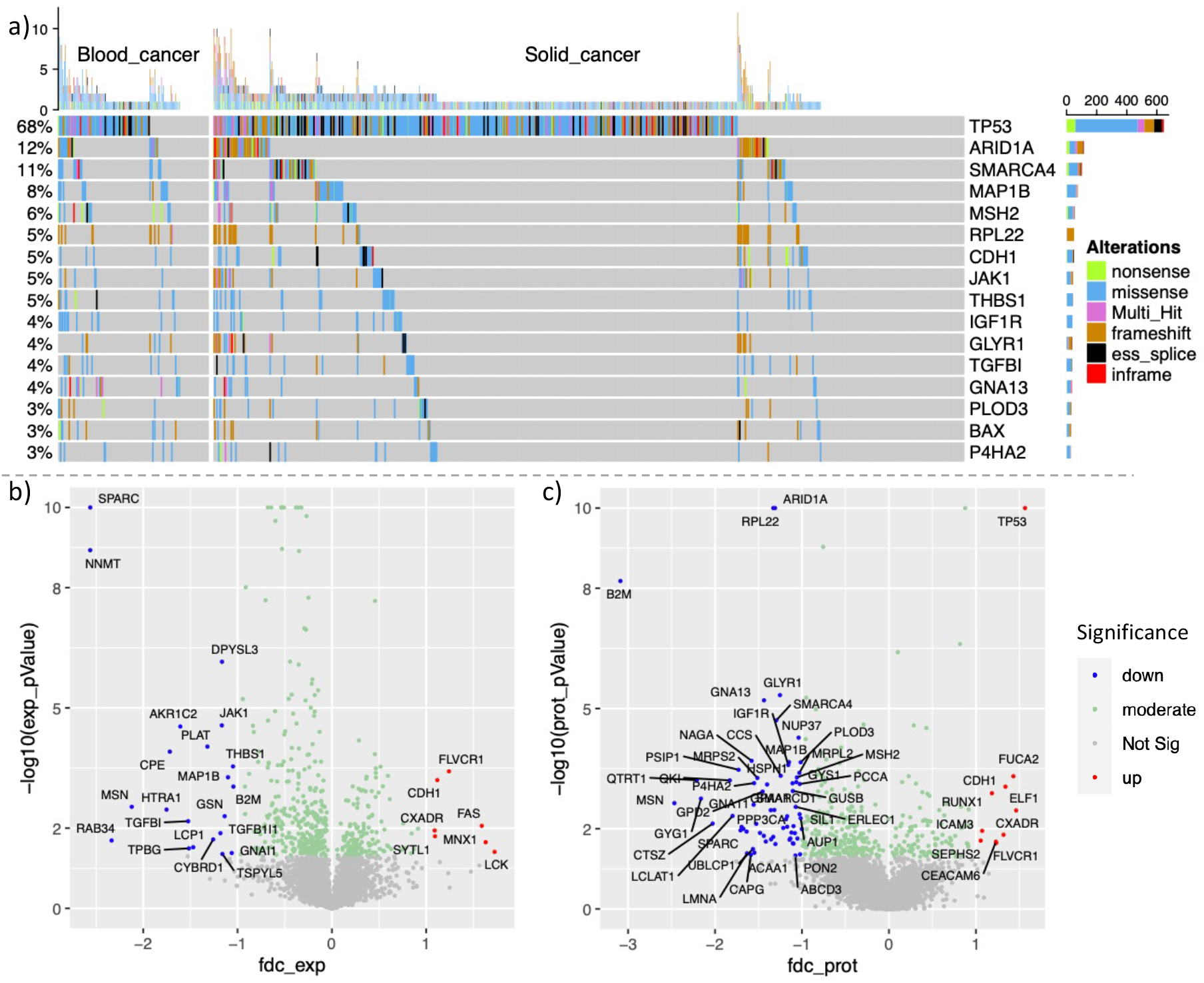
Genetic mutations and their effect on expression at the RNA and protein levels. a) The oncoplot of mutation landscape of genes that exhibited either up-regulated or down-regulated expression in either RNA or protein levels (by ComplexHeatmap^30^); only genes with a mutation frequency of at least 3% of whole cell line population were presented. b) The Volcano plot of differential expression RNAs. Each gene being compared must simultaneously have at least 10 mutated cell lines and 10 non-mutated cell lines. The x-axis indicates the log fold-change of RNA expression level, and the y-axis indicates the (negative) p-value of two-sided Wilcoxon rank sum test. c) The Volcano plot of differential expression proteins. We required that each gene being compared must simultaneously have at least 10 mutated cell lines and 10 non-mutated cell lines. The x-axis indicates the log fold-change of protein expression level and the y-axis indicates the (negative) p-value of two-sided Wilcoxon rank sum test.

#### Proteomics is an essential component in evaluating oncogenesis-related cancer gene mutations

We identified tens of differential expression genes (cell lines with gene mutation vs. wild type) that exhibited either up-regulated or down-regulated expression levels in RNA or protein levels. As presented in Fig. 3a, sixteen genes exhibited either up-regulated or down-regulated expression in either RNA or protein levels with gene mutation frequencies in or above 3% of the whole cell line population with genomic mutation profile. However, the oncogenesis impact as well as the clinical translation value for those genes are not well evaluated. As suggested in study^31^, gene mutations will affect the expression level and functionality (such as activating protein function in oncogenes^32^ and inactivating function in tumor suppressors^33^) of proteins. In order to evaluate the extent to which proteomic data can assist the identification of cancer genes (defined as mutated genes that causally implicated in oncogenesis^34^) and putative therapeutic target genes, we conducted a comparison for genes exhibiting differential expression (DEGs) at RNA level or DEGs at the protein level in terms of their overlap with cancer genes (1102 genes, last update 05/19/2023) that annotated in the expert-guided OncoKB database (https://www.oncokb.org/). The OncoKB is a precision oncology knowledge base and an FDA-recognized human variant database, which links cancer somatic molecular alterations with their tumor-specific clinical implications^24^.

In general, DEGs at the protein level obtained higher overlapping with OncoKB listed cancer genes (labelled as on-cokb_cancer_genes, Fig. 4). There are 18 genes that down-regulated at the protein level (labeled as down_protein_DEGs) that overlap with oncokb_cancer_genes (Fig. 4c), while only 4 genes that down-regulated at the RNA level (labeled as down_rna_DEGs, Fig. 4a) overlap with oncokb_cancer_genes. There was a similar overlap gene count between up-regulated genes at both RNA (labeled as up_rna_DEGs, Fig. 4b) and protein levels (labeled as up_protein_DEGs, Fig. 4d) with oncokb_cancer_genes. Next, we spotted the light on DEGs with mutation frequencies of 3% or higher across the cell lines. We observed 9 out of 13 DEGs at the protein level overlapped with oncokb_cancer_genes (Fig. 3a). Among them, two protein_mut_high_DEGs (ARID1A and TP53) are druggable genes annotated at the OncoKB database. The somatic alterations of tumor suppressor gene TP53 are one of the most common mutations in human cancers^35^, and the mutations of this gene were normally associated with poor prognosis in many cancers, such as head and neck cancer and breast cancer^36^. As annotated in OncoKB base, TP53 is a druggable gene for compound PC14586 for all solid cancer types that target TP53 alteration Y220C with level 3 clinical evidence ((https://www.oncokb.org/actionableGene). The compound PC14586 can selectively bind to p53 Y220C mutant protein and restores the transcriptional activity and conformation of p53 wild-type proteins, acting as a novel small molecule structural corrector^37^. Similar to one of the functional roles of TP53, gene ARID1A also acts as a tumor suppressor gene that prevents genomic instability^38^. Compounds PLX2853 and Tazemetostat are anti-cancer drugs that target the truncating mutations of gene ARID1A across all solid tumours with level 4 biological evidence (https://www.oncokb.org/actionableGenes). Between them, Tazemetostat was considered as a representative agent that target the histone methyltransferase enhancer Zeste homolog2^39^ (EZH2) in phase 2 clinical trails^40,41^ for ARID1A-deficient Ovarian Clear Cell Carcinoma^42^, which acts in a synthetic lethal manner in ARID1A-mutated cells. Our results demonstrated that proteomic data could help better identify protein coding gene biomarkers, which cannot be inferred solely based on RNA transcript levels, with the expressional and functional impact of genomic mutations on cancer genes related to cancer oncogenesis. The above findings further suggested that proteomic data is essential in evaluating oncogenesis-relate cancer genes with potential drug actionable alteration as anti-cancer drug targets.

**Figure 4.**
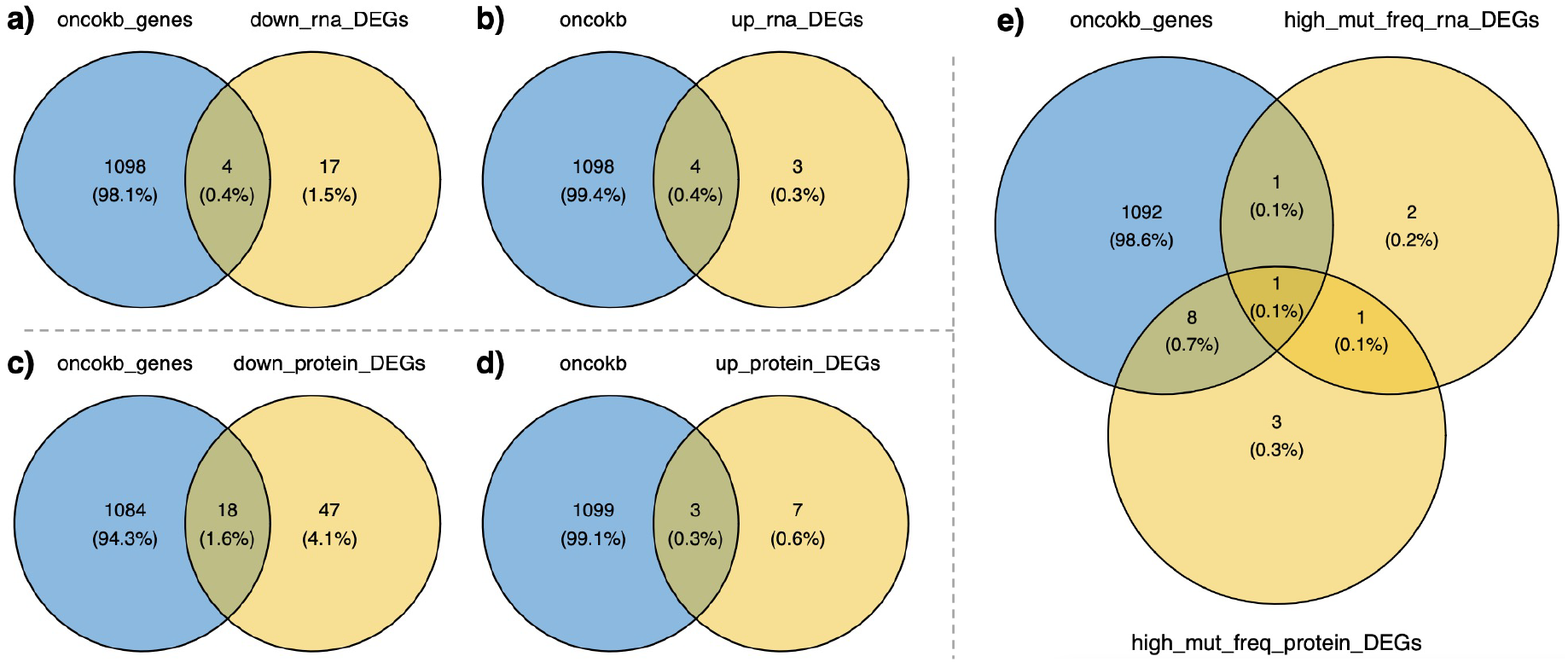
The association of differentially expressed genes (DEGs, either reflected in RNA or protein level) with OncoKB-annotated genes. a) Venn plot for the OncoKB-annotated genes and RNA down-regulated (gene-mutated vs. wild-type cell lines, same in the following subfigures b-d) genes. b) Venn plot for the OncoKB-annotated genes and RNA up-regulated genes. c) Venn plot for the OncoKB-annotated genes and protein down-regulated genes. d) Venn plot for the OncoKB annotated genes and protein up-regulated genes. e) Venn plot overview of overlapping frequency among RNA-DEGs, protein-DEGs, and OncoKB-annotated genes.

**Figure 5.**
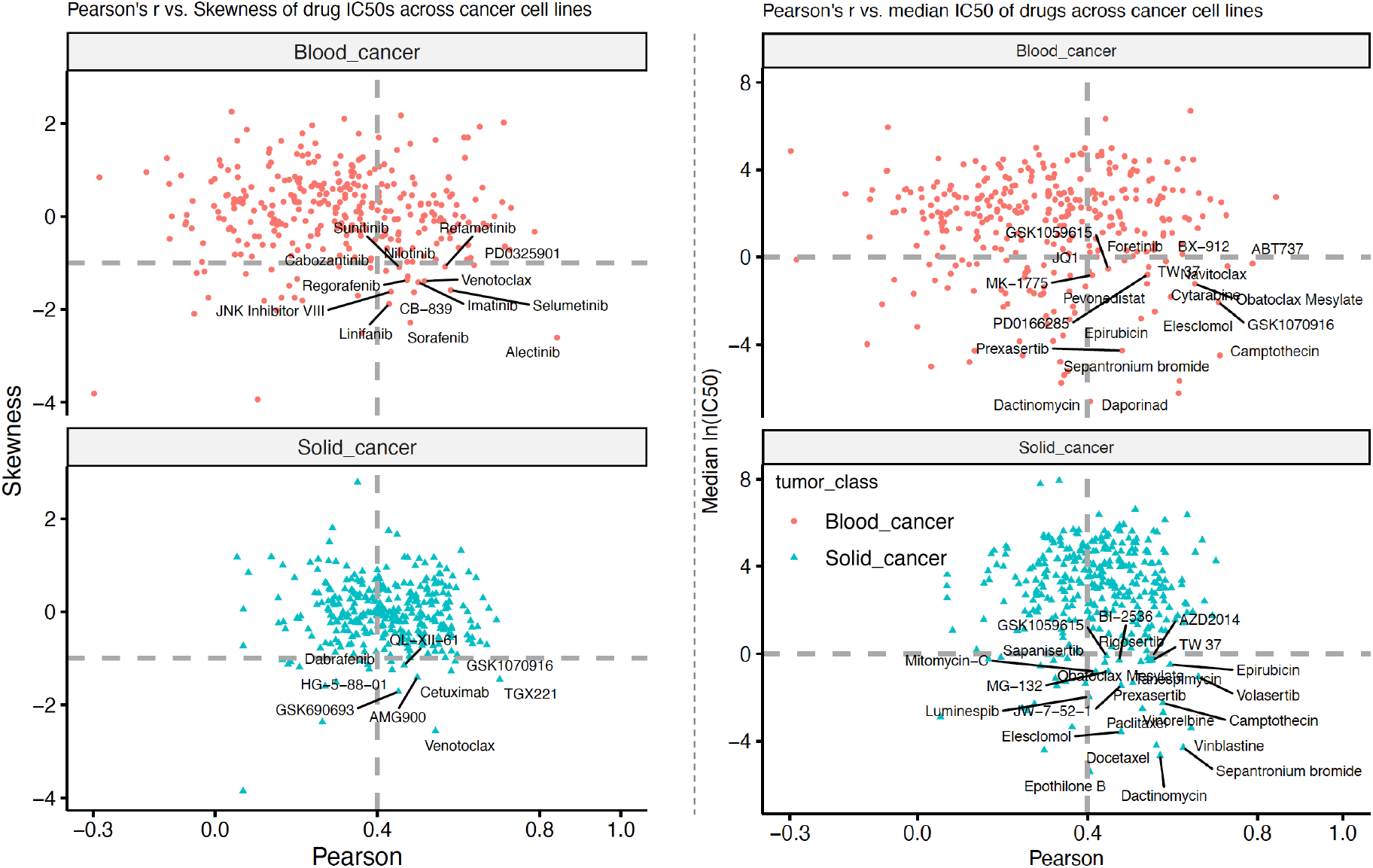
Predictive power and selectivity of drugs between hematological malignancies and solid cancers. a) Pearson’s r of true ln(IC50)s and predicted ln(IC50)s for drugs versus the skewness of ln(IC50)s across cancer cell lines. Blood cancers and Solid cancers were analyzed separately. b) Pearson’s r of true ln(IC50)s and predicted ln(IC50)s for drugs versus the median value of ln(IC50)s across cancer cell lines. Blood cancers and Solid cancers were analyzed separately.

### Predictive power and selectivity varied between hematological malignancies and solid cancers

In the analysis regarding high-confidence drug response prediction and drug selection for cancer cell lines in a previous study^16^, the authors did not describe the similarities and differences in the drug response predictions between hematological cancers and solid tumors. However, the difference between these two types of cancer contributes to the evident different RNA-protein correlations (Fig. 2) and different anti-cancer treatment prognosis^20^, suggesting that it is vital to discuss them separately.

Moreover, we noticed that the author adopted Fisher-Pearson coefficient skewness of ln(IC50) values as a criterion for the prioritization of candidate promising anti-cancer drugs^16^. However, this strategy may conceal some drugs that have low skewness coefficients. More importantly, the concealed drugs also have low ln(IC50) values, whereas low IC50 value is an informative indicator for screening candidate drugs in drug cell viability assays^43^. Therefore, we followed the drug prioritization approach^16^ and reanalyzed the predicted results from our bi-modal model by separately analyzing cell lines of hematological malignancies and solid tumors. We compared the selected drugs between the analysis results using Fisher-Pearson coefficients and ln(IC50) skewness as the drug sensitivity prioritizing metric and the result that replacing skewness with median ln(IC50) value of each compound as the drug sensitivity prioritizing screening criteria. To obtain high-confidence Pearson correlation statistical results, we excluded drugs that were assayed in less than 15 cancer cell lines for hematological malignancies and solid tumors, respectively.

As expected, adopting the median ln(IC50) values of drugs as the screening criterion can prevent many anti-cancer compounds from being excluded when adopting skewness of ln(IC50)s as the drug prioritizing criterion (Fig. 5). We also noticed that the Pearson coefficients for all drugs in solid cancer are higher than zero, while a few drugs show poor Pearson coefficients in hematological malignancies (Fig. 5b). This may be due to the fact that there are fewer *in vitro* drug response experiments (cell viability assays) conducted on hematological malignancy cell lines in the GDSC database than on solid tumor cell lines, making it difficult for the model to learn and predict the drug response on unknown data from blood cancer cell lines in the test set. Due to the same reason, we obtained slightly fewer high-confidence candidate drugs under the threshold of Median ln(IC50)<0 and Pearson coefficient > 0.4 in hematological malignancies than that observed in solid cancers (Fig. 5b). Moreover, we have also observed many differences for the drugs that are commonly practiced for both hematological malignancies and solid tumors which presenting different treatment efficacy and predictive power. For instance, the median ln(IC50) for drug Daporinad obtains a Pearson score (ground-truth ln(IC50) vs. predicted ln(IC50)) higher than 0.61 among hematological malignancies with a median ln(IC50) of -6.23, while only exhibited a person’s r of 0.36 in solid cancer cell lines (Fig. 5b). Daporinad is a small molecule with potential capability of anti-cancer and anti-angiogenesis, which could inhibit the production of vascular endothelial growth factor (VEGF) in cancer cells (https://www.cancer.gov/publications/dictionaries/cancer-drug/def/daporinad). It is also act as a nicotinamide phosphoribosyltransferase (NAMPT) inhibitor^44^, which was proven to be effective in some chronic myeloid leukemia patients^45^. Such results cannot be observed under the combination metrics of Pearson score (ground-truth ln(IC50) vs. predicted ln(IC50)) and skewness of ln(IC50) values. Our results demonstrated that using the median ln(IC50) of drugs as a criterion could better prioritize anti-cancer compounds than skewness of ln(IC50) values, and the predictive power and selectivity varied between hematological malignancies and solid cancers. Consequently, analyzing drug responses for these two cancer types separately could help better identify cancer-type-specific promising anti-cancer drugs.

## Discussion

Based on the fact that the clinical trials are very costly and difficult to evaluate the *in vivo* drug response sensitivities of multiple anti-cancer drugs in parallel, the cancer cell lines still remain as the most commonly used *in vitro* model for anti-cancer drug treatment studies. Unfortunately, due to the limited protein quantification data, previous pharmacogenomic studies are mainly based on other omics data, such as genomic mutations and RNA expression profiles. However, none of them can actually infer the protein quantification levels, as reflected by the moderate or low pan-cancer gene-wise and cell-line-wise RNA-protein expression correlation coefficients (Fig. 2). In practice, proteins are the basic functional units of cells and the direct drug targets for most anti-cancer medicines^14^. More specifically, the chemical structures of anti-cancer drugs control the circuitries of the complex cellular networks and to modulate protein interactions and activities^15^. Therefore, it is necessary to bridge the gap in our understanding of pharmacodynamics by integrating protein profiles. Fortunately, the recently published large-scale ProCan-DepMapSanger proteomics map, which quantifies more than 8,000 protein-coding genes, enables us to conduct pharmacogenomic studies in cell lines at a pan-cancer scale^16^. Our proposed bi-modal drug response prediction neural network BDRN demonstrated that protein data could achieve comparable and slightly better performance in drug response prediction task than that in RNA sequencing data across cancer cell lines. The integration of drug chemical feature outperforms such task (DeepOmicNet^16^) conducted solely based on a single type of biological omics data. Additionally, integrating the chemical features of drug molecules (represented as graphs) enhances the scalability of BDRN, enabling it to predict drug molecules that were not encountered during the training process.

Diversities between hematological cancers and solid tumors are also reflected at cell-line-wise RNA-protein expression correlation coefficients and emerge in the differences in anti-cancer drug responses. Based on our results (Fig. 2a-b), the RNA-protein expression correlations among hematological cancers are significantly higher than those observed among solid cancer cell lines. Since proteins are the cellular functional units and the therapeutic target of many drugs, we believe there exists lot of drug treatment heterogeneity between these two cancer classes. We systematically compared the drug response sensitivities and ML prediction results; drug response heterogeneity was observed in many drugs. For instance, drug Daporinad was benchmarked with a Pearson score higher than 0.6 with a median ln(IC50) less than -6 in hematological cancers, while obtained a Pearson score lower than 0.37 in solid cancer cell lines. Daporinad has been used in several clinical trials for the exploration of anti-cancer therapy for many cancer types, such as Melanoma (a kind of skin cancer), as well as other hematological malignancies such as Cutaneous T-cell Lymphoma and B-cell Chronic Lymphocytic Leukemia (https://go.drugbank.com/drugs/DB12731). This result further supports the well-known view that the anti-cancer drug responses may differ between hematological and solid cancers. Therefore, drug response experiments and treatment recommendations should be considered separately between these two cancer classes.

The drawback of a large proportion of missing values in protein quantification profiles could be remitted by referring to the expression features of RNA transcripts. The characteristics of protein molecules still hinder the quantification of their expression levels more challenging and costly than sequencing RNA transcripts. Due to both technical and biological aspects^16^, the DIA-MS-based proteomic data contain a large proportion of missing values. The data processing methods for protein missing values may greatly influence the analytical results. We reasoned that this drawback of protein missing data to some extent could be remitted by referring to the data distribution characteristics of RNA transcripts. To better characterize these protein data, we compared the results of the differential expression analysis and RNA-protein correlation analysis when missing values were deleted (in such a scenario, the corresponding RNA data of genes with missing protein values were also ignored during analysis) or when missing values were replaced as zeros. We found that replacing missing values as zeros can better retain the molecular expression features in both RNA and protein data, especially those dropout genes with low expression levels of both RNAs and proteins.

The exploration of potential biomarkers plays a crucial role in advancing our understanding of tumor development and identifying potential targets for drug intervention. A greater number of differentially expressed biomarkers were identified at the protein level as compared to the RNA level (Fig. 3b-c). Since proteins are the primary therapeutic targets for most anti-cancer drugs, we reasoned that the identified protein biomarkers could be integrated with public expert-annotated precision oncology knowledge databases such as OncoKB base to evaluate the potential clinical translation value. By carefully comparing the cancer genes and druggable genes that were annotated in OncoKB base, we identified more carcinogenesis-related genes and promising druggable genes based on proteomic data of cancer cell lines compared to transcriptomic data (Fig. 4). To the best of our knowledge, our method introduces a novel perspective for comparing differentially expressed biomarkers between transcriptomic and proteomic data, offering original insights into analyzing these data types together.

Above all, proteomics plays a vital role in cancer characterization, offering profound insights and opportunities for advancing our understanding and cancer treatment. Previously, the characteristics of protein molecules made the quantification of their expression levels more difficult than sequencing RNA transcripts, resulting in many pharmacogenomic analyses on large-scale pan-cancer cell lines with other omics data types such as genomics and transcriptomics. Fortunately, more and more protein sequencing techniques are emerging, such as protein sequencing at single cell resolution^46^. It is now necessary to fill the gap in the central dogma of genetics^23^ with the enlargement of protein data. Therefore, there is a pressing need to evaluate the inference capability of drug response and cancer vulnerability based on the proteomic data on a larger scale. By systematically characterizing protein features over 8,000 proteins across 949 cancer cell lines^16^, our results offer a reference for the proteomics analysis of in the context of cancer drug response modeling. Overall, proteomic data could provide us with new insights into perturbational biology, which differs from transcriptomic and genomic data types^16^, enabling us to identify putative therapeutic targets for future studies^8^.

## Methods

### Data availability

#### Raw data collection

All raw data used in this study are publicly available. The complete processed map of genomic mutations of GDSC database cancer cell lines was downloaded in supplementary Table S2C (CellLineVariants) from study^17^. The RMA normalized RNA expression data of GDSC cancer cell lines were processed and can be downloaded from https://ibm.ent.box.com/v/paccmann-pytoda-data/folder/127994700682^4^. The processed protein measurements of the ProCan-DepMapSanger dataset can be found in the subsheet “Prot matrix excl single-peptide” of supplementary Table S2, which was filtered and quality controlled from previous study^16^. The processed protein data of the ProCan-DepMapSanger dataset is also available at https://zenodo.org/record/6563157 and https://github.com/EmanuelGoncalves/cancer_proteomics^16^. The list of oncogenesis-related cancer genes and expert-guided druggable genes was downloaded from OncoKB database (https://www.oncokb.org/, last update 05/19/2023 when downloading), an FDA-recognized human variant knowledge base for precision oncology.

### Data Preprocessing

#### ML training/test set splitting

For the machine learning procedure, we used the previously divided training/testing cell line list that was conducted in study^16^, in which 80% of cancer cell lines were selected as training set for the 5-fold cross-validation (random seed = 42) and hyper-parameter selection, while the remaining 20% of cancer cell lines were selected as independent test set. Two additional filtering steps were also conducted before the training of our ML model. There are two versions of GDSC datasets (GDSC1 and GDSC2). As described in the documentation section of the GDSC website (https://www.cancerrxgene.org/help#t_curve), there are many Cell viability assays and experiments have been repeated with improved experiment equipment and procedures, and it is recommended to use the results of GDSC2 dataset when duplicate IC50 values exist. In addition, the inclusion of duplicate drug screening results may introduce sample-size bias during the machine learning training and testing processes. Therefore, we followed the suggestion from GDSC documentation and removed the duplicate results from the GDSC1 dataset. Moreover, only cancer cell lines with transcriptomic and proteomic profiles were considered for the ease of comparison, resulting a total of 911 cancer cell lines were retained in our analysis. Among them, there are 761 solid cancer cell lines and 150 hematological malignancy cell lines.

#### Protein missing value processing

When performing RNA-protein correlation analysis, we used two different methods to handle proteins with missing values. One method involved removing the missing data, which also requires ignoring RNA expression data for genes with missing protein quantification values. The other method involved replacing all missing protein values with zeros, allowing us to retain all RNA expression data.

#### Drug chemical feature generation

To generate drug molecular graph features, we first downloaded the drug SMILES (Simplified Molecular Input Line Entry System) format files from https://ibm.ent.box.com/v/paccmann-pytoda-data/folder/91701932285^4^, which consists SMILES string from GDSC^2^, CCLE^3^, and ChEMBL^47^ databases. The redundant entries were removed. The drug molecular chemical information was generated via *Chem*.*MolFromSmiles()* function of rdkit package^26^ (https://www.rdkit.org/). The drug feature was generated via the function of *ConvMolFeaturizer()* from package deepchem^25^ (https://github.com/deepchem/deepchem/). A total of 334 drugs with drug molecular chemical features (represented as drug graphs) were included in our study.

### Workflow Overview of BDRN

#### Feature extractors for transcriptomic and proteomic data

The feature extractors for protein and RNA data are both composed of MLPs (multilayer perceptrons, denoted as *M*), and the feature extractor for drug graph is GNN (graph neural network, denoted as *G*). We denote the protein data and RNA data as *XP, XR*, and the corresponding latent vector representation mappings are:

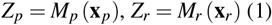

and the drug graph input as *XD*, and the corresponding latent vector representation mapping for drug graph is:

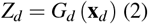

#### Drug response predictor

Drug response predictor (denote as *P*) is a three-layer MLPs with a dropout rate of 0.5, *pd* represents the latent vector of proteomics data concatenates drug graph latent vector, and *rd* represents the latent vector of proteomics data concatenates drug graph feature latent vector. We denote the output as:

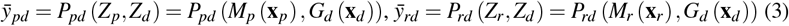

#### Loss function

The performance of BDRN framework and the drug response predictor was evaluated based on Mean squared error (MSE) loss:

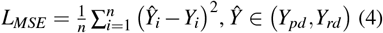

#### Hyper-parameters

We introduced two hyper-parameters. The hyper-parameter *h*_*dim* is set to control the number of nodes for the first layer of proteomic/transcriptomic feature extractor, and the hyper-parameter *o*_*dim* is set to control the number of nodes for the omic latent vector (Fig. 1). We also rerun the cross-validation for baseline model DeepOmicNet, which comprises two hyper-parameters *hidden*_*width* to control the number of hidden layer neurons and hyper-parameter *group* to control the number of grouped bottlenecks^16^. The best hyper-parameters based the loss of cross-validation were tabularized in Table S1 and Table S2 for baseline model DeepOmicNet and our BDRN frameworks.

#### Performance metric

We conducted a five-fold cross-validation (CV) for hyper-parameter selection to reduce data splitting bias during the training process. We averaged the predicted ln_IC50s on the test set using the 5-fold models (the model that obtained the minimal validation loss of each fold) to obtained a reliable prediction. After obtaining the averaged ln_IC50s of each drug-cell_line pair, we calculated the cell_line-wise Pearson’s r for each drug individually. The median and mean of the cell_line-wise Pearson’s r across all drugs were considered as the metrics for model performance comparison.

#### Statistical Analysis

All statistical testings were performed in R 3.6.3 environment by two-sided Wilcoxon rank-sum test, and the significant level was set to p<0.05 unless specified.

## Acknowledgements

This research was substantially sponsored by the research project (Grant No. 32170654 and Grant No. 32000464) supported by the National Natural Science Foundation of China and was substantially supported by the Shenzhen Research Institute, City University of Hong Kong. The work described in this paper was substantially supported by the grant from the Research Grants Council of the Hong Kong Special Administrative Region [CityU 11203723]. The work described in this paper was partially supported by the grants from City University of Hong Kong (2021SIRG036, CityU 9667265, CityU 11203221) and Innovation and Technology Commission (ITB/FBL/9037/22/S).

