## Supplementary file for "Drug Response Modeling across Cancers: Proteomics vs. Transcriptomics"

**supplementary figures**

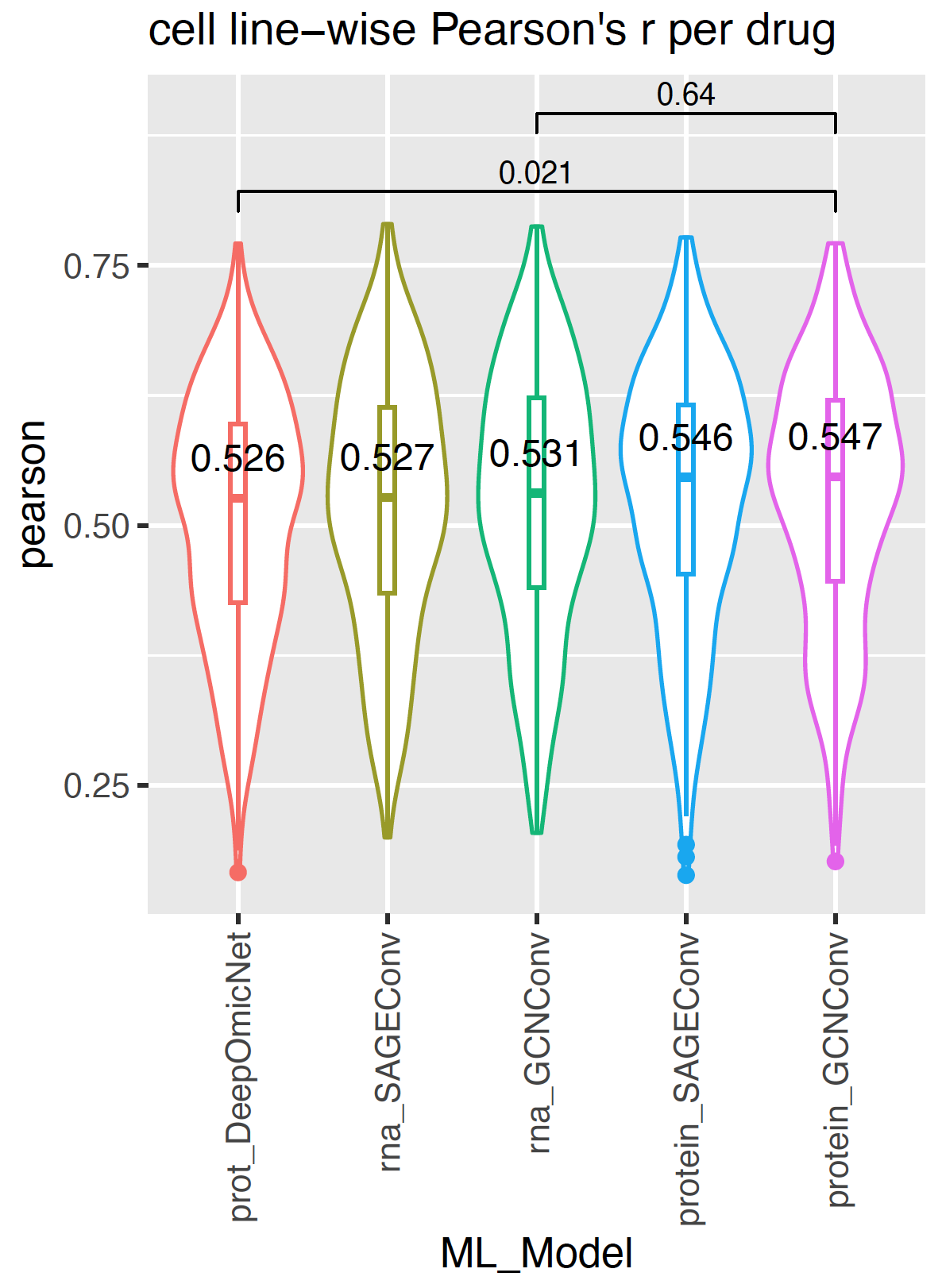

**Fig. S1.**

The performance of GNN models compared to the baseline model DeepOmicNet, the median and mean Pearson’s r per drug across cell lines were applied as performance metrics, referring to the previous study.

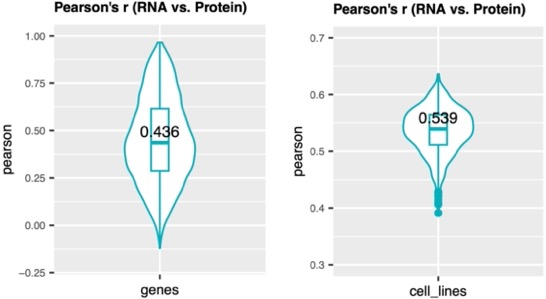

**Fig. S2.**

Pan-cancer cell line RNA-protein correlations. a) Boxplot with violin plot for the gene-wise correlation between RNA and protein expression levels; missing values of proteins were replaced as zeros. b) Boxplot with violin plot for the cell-line-wise correlation between RNA and protein expression levels; missing values of proteins were replaced as zeros.

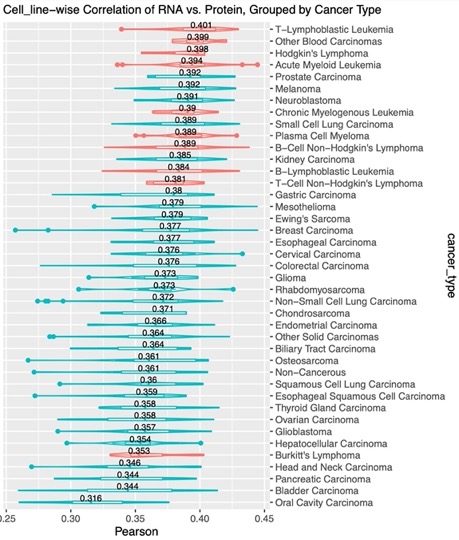
**Fig. S3.**

Boxplot with violin plot for the cell line wise correlation between RNA and protein expression level, missing values of proteins were ignored during the correlation analysis, and the corresponding RNA values for genes with missing protein values were also ignored.

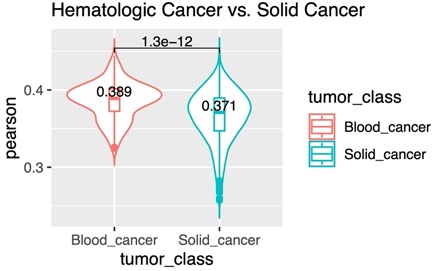
**Fig. S4.**

Boxplot for the comparison of RNA-protein cell-line-wise correlation coefficients between hematological cancers and solid cancers, missing values of proteins were ignored during and the correlation analysis, the corresponding RNA values for genes with missing protein values were also ignored.

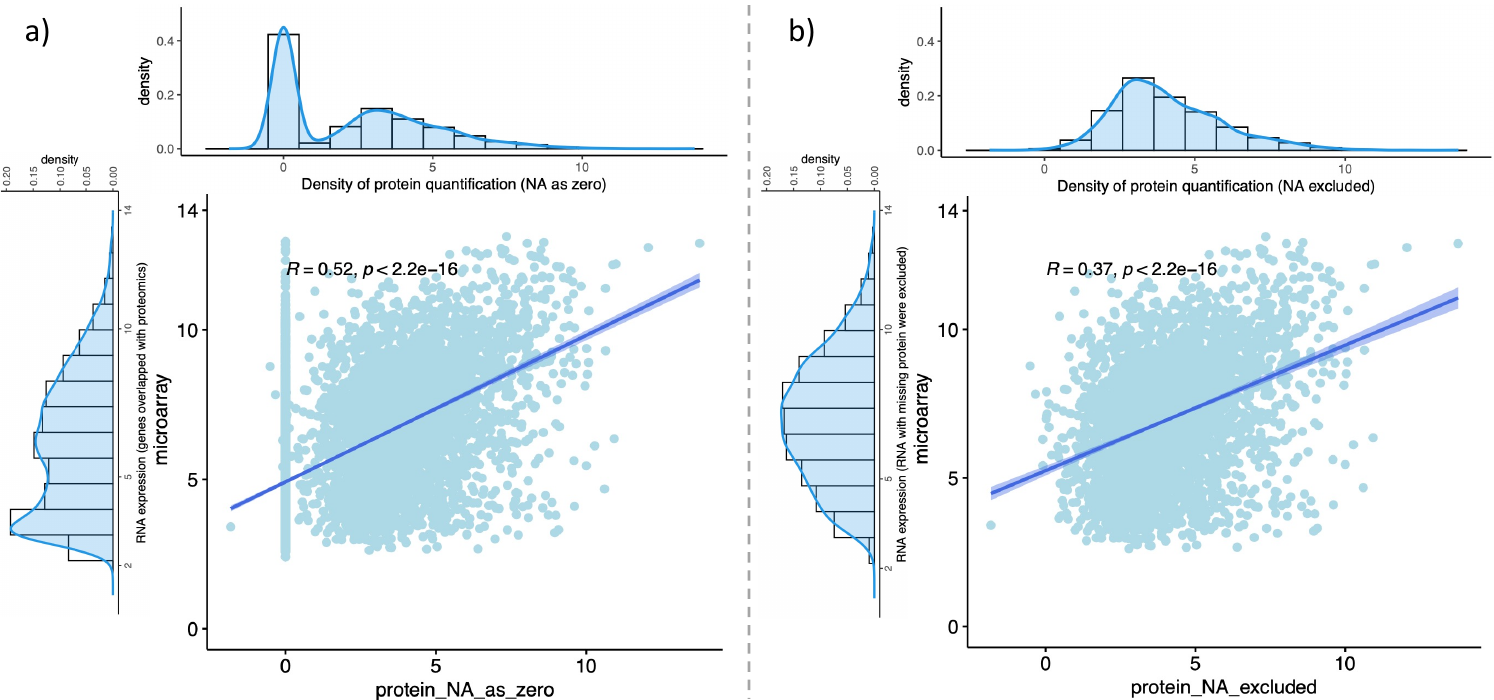
**Fig. S5.**

The mRNA-protein correlation scatter plot for glioblastoma cell line GAMG. a) scatter plot for RNA and corresponding protein expression values, where missing values of proteins were replaced as zeros. b) scatter plot for mRNA values and corresponding protein values, where both the RNA values and the corresponding proteins of genes with missing protein values were ignored.

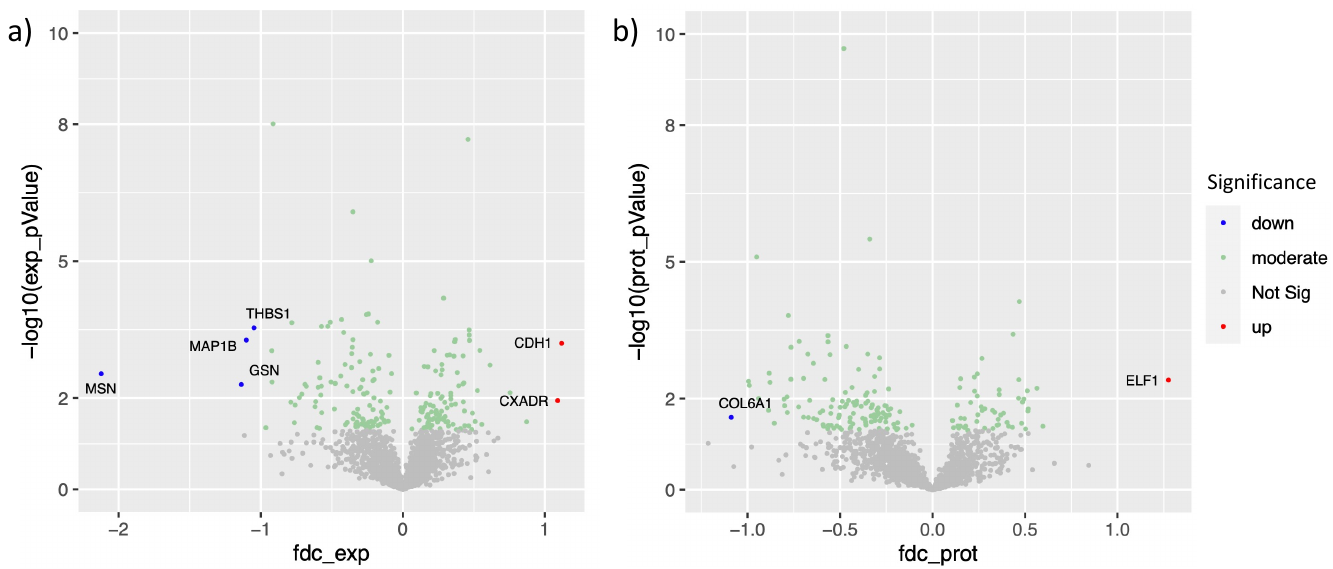
**Fig. S6.**

The impact of genetic mutation at RNA and protein expression levels, which both the RNA values and the corresponding proteins of genes with missing protein values were ignored during differential expression analysis. a) The Volcano plot of differential expression RNAs. Each gene being compared must simultaneously have at least 10 mutated cell lines and 10 non-mutated cell lines. The x-axis indicates the log fold-change of RNA expression level, and the y-axis indicates the (negative) p-value of the two-sided Wilcoxon rank sum test. b) The Volcano plot of differential expression proteins. Each gene being compared must have at least 10 mutated cell lines and 10 non-mutated cell lines simultaneously. The x-axis shows the log fold-change of RNA expression level, and the y-axis indicates the (negative) p-value of the two-sided Wilcoxon rank sum test.

**supplementary tables**

supplementary Table 1

| data_type | model | hidden_width | group |
| --- | --- | --- | --- |
| protein | DeepOmicNet (baseline) | 3000 | 1500 |

supplementary Table 2

| data_type | Conv_layers | h_dim | o_dim |
| --- | --- | --- | --- |
| RNA | BDRN (GCNConv) | 1024 | 1024 |
| RNA | BDRN (SAGEConv) | 1024 | 512 |
| Protein | BDRN (GCNConv) | 1024 | 1024 |
| Protein | BDRN (SAGEConv) | 1024 | 1024 |

The hyper-parameters for baseline model DeepOmitNet are ([3000,3000], [3000,1500], [3000,1000], [3000,500], [1500,1500], [1500,500], [500,500]). The hyper-parameters for BDRN are ([1024,1024], [1024,512], [1024,256], [512,512], [512,256], [256,256] for both transcriptomic and proteomic data. The best hyper-parameters for baseline model and our BDRN models with minimum 5-folds cross validation losses are summarized in Table S1 and Table S2, respectively.
